# Sequence Generation and Phylogenetic Inference with Generative Flow Networks

**DOI:** 10.64898/2026.04.08.717239

**Authors:** Qichen Huang, Carlos Michel Mourra-Diaz, Xiaozhen Wen, David Payette, Archer Y. Yang

**Affiliations:** Mila – Québec AI Institute; McGill University; Université de Montréal; Department of Mathematics and Statistics, McGill University Montréal, QC, Canada

## Abstract

Phylogenetic inference remains computationally challenging due to the exponentially growing tree topology search space, and current methods rely heavily on multiple sequence alignments (MSAs) which are expensive and error-prone. We propose AncestorGFN, a proof-of-concept approach leveraging Generative Flow Networks (GFlowNets) for simultaneous sequence generation and phylogenetic exploration without requiring explicit MSAs. Our method learns to generate sequences matching a target distribution while the flow trajectories implicitly encode structural relationships among sequences. We demonstrate that greedy traceback on maximum-flow trajectories recovers shared intermediate states suggestive of common ancestry, and evaluate on the let-7 microRNA family where the learned flow structure qualitatively captures phylogenetic branching patterns. Furthermore, beam search at inference time discovers novel sequences clustering near known targets, suggesting applications in *de novo* sequence design. This work establishes an initial foundation for alignment-free phylogenetic exploration using generative models.

## 1 Introduction

Phylogenetic inference, which reconstructs evolutionary relationships from molecular sequences, remains computationally challenging. The number of possible tree topologies grows exponentially with the number of taxa: there are (2*n* − 5)!! unique unrooted bifurcating tree topologies on *n* species (Zhou et al., 2024). Standard methods including parsimony, maximum-likelihood, and Bayesian approaches have been developed to explore this vast space (Haber & Velasco, 2024), but these rely on multiple sequence alignments (MSAs), which are computationally expensive and can introduce errors that propagate to the inferred trees.

Generative Flow Networks (GFlowNets) offer a promising alternative for exploring large discrete spaces (Bengio et al., 2021; 2023). GFlowNets learn to sample objects with probability *P* (*x*) ∝ *R*(*x*), where *R*(*x*) is a reward function, with theoretical connections to variational inference (Malkin et al., 2023) and demonstrated effectiveness in discrete optimization (Zhang et al., 2023; Pan et al., 2022). Structurally, GFlowNets can be viewed as directed acyclic graphs (DAGs) where nodes represent states, edges denote transitions, and flow corresponds to trajectory probabilities (Figure 1).

**Figure 1:**
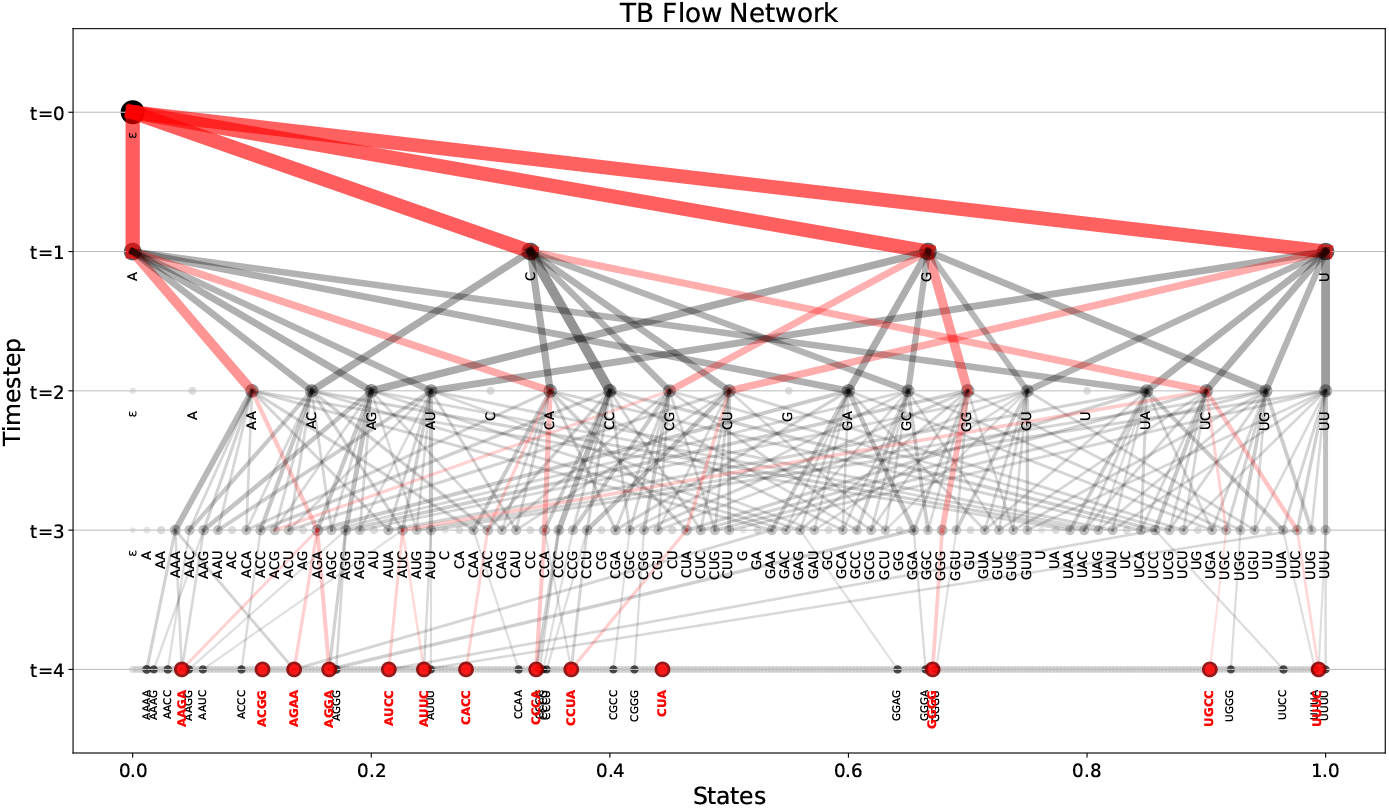
Schematic overview of a GFlowNet trained with the Trajectory Balance (TB) objective. Target sequences (red) represent the desired distribution; black nodes are off-target candidates. Trajectories show sampled paths through the flow network, with maximum-flow traceback paths highlighted in red.

Generative models have been applied to both sequence generation and phylogenetic inference. Jain et al. (2022) demonstrated GFlowNets for *de novo* protein and DNA sequence generation. PhyloGFN (Zhou et al., 2024) applies GFlowNets to tree topology generation, while PhyloGAN (Smith & Hahn, 2023) uses GANs for generating tree topologies and branch lengths. Phylo-Diffusion (Khurana et al., 2024) conditions diffusion models with phylogenetic knowledge for species image generation. However, using generative models to simultaneously generate sequences and explore phylogenetic relationships from the generation trajectories remains largely unexplored.

Here, we propose AncestorGFN, which leverages GFlowNets for both sequence generation and phylogenetic exploration without requiring explicit MSAs. Our primary contributions are: (1) reframing GFlowNet flow trajectories as a lens for qualitative phylogenetic analysis, where shared intermediate states suggest common ancestry; (2) demonstrating that FL-DB with carefully designed intermediate rewards enables effective exploration of large sequence spaces; and (3) showing that beam search at inference time can propose novel candidate sequences near known functional targets—bridging generative modeling with *de novo* sequence design.

## 2 Method

AncestorGFN adapts GFlowNets to both sequence generation and phylogenetic inference. We generate RNA sequences matching a target distribution and infer phylogenetic relationships from the learned flow network.

### 2.1 GFlowNet Training Objectives

Our implementation mainly uses the Forward-Looking Detailed Balance (FL-DB) training objective (Pan et al., 2023), which addresses challenges of long trajectories and sparse rewards by incorporating intermediate reward signals.

#### State Space

Each state represents a sequence. The initial state is an empty sequence *ϵ*, and terminal states are complete sequences. States track the time step *t* to maintain the DAG property.

#### Action Space

We define three action types:

- **Insertions:** Insert a nucleotide (A, U, G, C) at any valid position
- **Substitutions:** Replace any existing nucleotide with a different nucleotide
- **Deletions:** Remove a nucleotide from any valid position

#### Reward Function

We implement several intermediate reward strategies based on sequence similarity to targets (see Appendix A for algorithmic details).

#### Training Objectives

We compare three GFlowNet training objectives:

*Trajectory Balance (TB)* (Malkin et al., 2022): *For a complete trajectory τ* = (*s*_0_ → *s*_1_ → · · · → *s*_*n*_ = *x*):

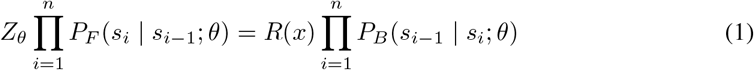

where *Z*_*θ*_ is a learnable partition function estimate.

*Detailed Balance (DB)* (Bengio et al., 2023): *For each transition s* → *s*′:

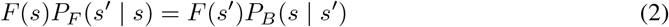

where *F* (*s*) is the learned state flow function, with boundary condition *F* (*x*) = *R*(*x*) at terminal states.

*Forward-Looking Detailed Balance (FL-DB)* (Pan et al., 2023): The key insight is to reparameterize the flow function to incorporate intermediate energies. Given an energy function *E*(*s*) defined on all states (not just terminal), we define the **transition energy**:

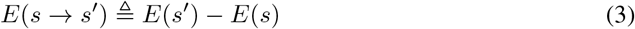

and the **forward-looking flow**:

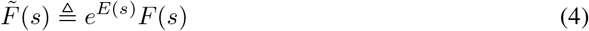

This reparameterization factors out the energy already accrued at state *s*, so 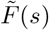 only depends on future transitions. Equivalently, 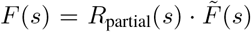, where *R*_partial_(*s*) = *e*^−*E*(*s*)^ is the partial reward at state *s*. Substituting into Eq. 2 yields the **FL-DB constraint**:

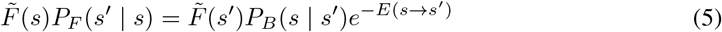

The corresponding loss function in log-space is:

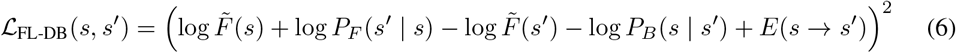

In our implementation, we define the intermediate energy *E*(*s*) using sequence similarity to targets, such that *R*_partial_(*s*) = *e*^−*E*(*s*)^ provides a partial reward signal at each step. This enables more efficient credit assignment and faster convergence, especially for long sequences.

### 2.2 Flow Traceback for Phylogeny Inference

To infer phylogeny from the trained GFlowNet, we leverage the flow structure of the DAG. We first compute edge flows by forward-propagating from the source: for each edge (*s* → *s*′), the flow is defined as:

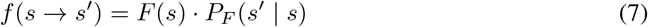

where *F* (*s*) is the cumulative flow at node *s*.

We then apply **greedy backtracking** to reconstruct evolutionary trajectories: starting from each target terminal state *x*, we iteratively select the parent node with maximum incoming flow:

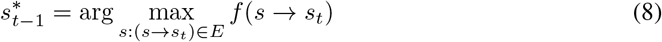

This process continues until reaching the root (empty sequence). The intersection points of these trajectories correspond to shared intermediate states that we interpret as putative common ancestors under the generative model.

## 3 Experiments

### 3.1 Case Study 1: Short RNA Sequences (4BP)

We evaluate AncestorGFN on short RNA sequence generation using an alphabet of four nucleotides {*A, U, G, C*} with maximum sequence length of 4 and 5 discrete time steps. We define 13 target RNA motifs as the desired targets as an example, including sequences such as AUUC, AUCC, CACC, CUA, CCUA, CCCA, GGGG, and others with varying sequence complexity.

#### Training Objective Comparison

We compare three GFlowNet training objectives: Trajectory Balance (TB) (Malkin et al., 2022), Detailed Balance (DB) (Bengio et al., 2023), and Forward-Looking Detailed Balance (FL-DB) (Pan et al., 2023). For TB and DB, we use a simple sparse reward signal that returns 1.0 for exact target matches and 0.1 otherwise. For FL-DB, we employ AlignmentReward (Appendix A.1) which provides partial credit for sequences similar to targets, enabling better credit assignment during training.

All models are trained for 20,000 episodes with learning rate 3 × 10^−3^. Figure 2A shows the training loss comparison. TB, which only receives reward at terminal states, suffers from poor credit assignment on longer trajectories. In contrast, DB and FL-DB converge faster by learning per-state flow estimates that provide more localized gradient signals (Pan et al., 2023). Figure 2B shows normalized convergence dynamics, demonstrating that all three objectives successfully learn the target distribution. Notably, we find FL-DB achieves higher mean reward than TB and DB, as its partial reward signal provides better guidance during exploration. A detailed comparison of training objectives is provided in Appendix B.

**Figure 2:**
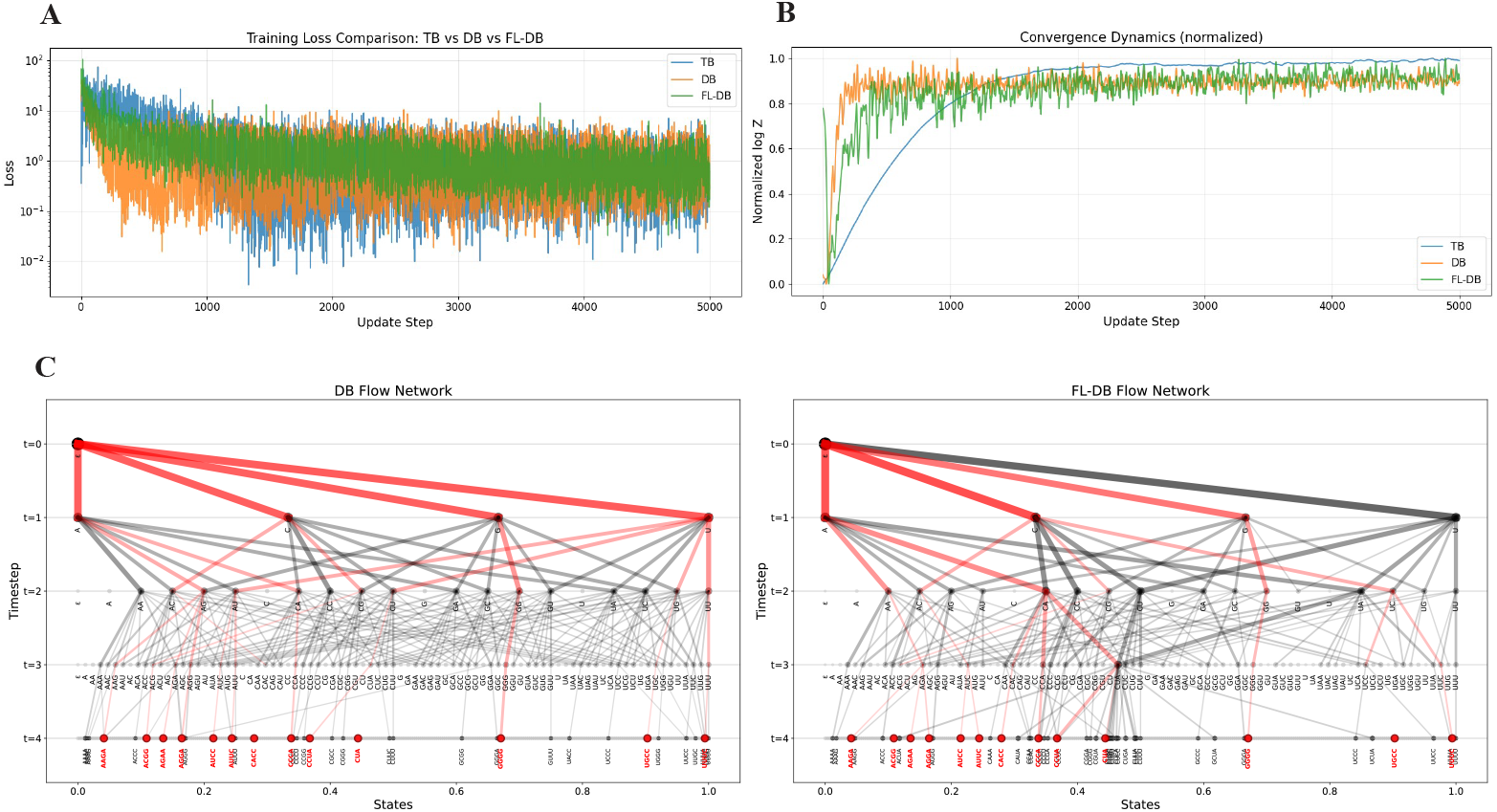
Training comparison of GFlowNet objectives on short RNA sequences. (A) Training loss curves for TB, DB, and FL-DB objectives. (B) Normalized partition function estimates showing convergence dynamics. (C) Flow network visualization for DB (left) and FL-DB (right) models, with target sequences highlighted in red and maximum-flow trajectories shown. The TB flow network is shown in Figure 1.

#### Flow Network and Phylogeny Inference

The trained flow networks reveal phylogenetic structure through their sampled trajectories. Figure 1 shows the TB model’s flow network, while Figure 2C displays the DB and FL-DB networks. Applying greedy traceback, we find that some sequences share common ancestors across different training objectives. For example, both CCCA and GGGG pass through a shared ancestral state, while AAGA and AGGA both trace back to the same ancestral sequence AGA. These branching patterns suggest that the GFlowNet’s learned flow structure captures relationships among sequences that are qualitatively consistent with evolutionary intuition. Furthermore, because the learned DAG structure in GFlowNets allows for multiple parent nodes, some target sequences are inferred to have more than one ancestral predecessor.

### 3.2 Case Study 2: Long Sequences and Let-7 MicroRNA (10BP)

We evaluate AncestorGFN’s scalability by extending to 10bp RNA sequences, where the state space grows to 4^10^ = 1, 048, 576 possible sequences—over 4,000× larger than the 4bp case.

#### TB vs FL-DB on 10bp Sequences

We first compare TB and FL-DB on 100 randomly generated 10bp target sequences. For FL-DB, we explore different partial reward schemes: (1) basic HammingReward providing normalized edit similarity, (2) EntropyWeightedHammingReward (Appendix A.2) which boosts rewards for complex sequences to prevent mode collapse on simple repetitive patterns, and (3) AdaptiveHammingReward (Appendix A.3) which dynamically decays rewards for frequently hit targets.

After 102,400 training episodes, FL-DB significantly outperforms TB in target coverage: FL-DB discovers 10/100 unique targets cumulatively while TB finds only 2/100. This demonstrates that partial reward signals become increasingly important as the search space grows, providing gradient information even when exact target matches are rare.

#### Let-7 MicroRNA Family

We evaluate on biologically relevant sequences: the let-7 microRNA family, one of the most conserved miRNA families across species (Roush & Slack, 2008). We extract 612 let-7 sequences from 107 species and select positions 10–19 (a 10bp variable region identified via sliding window with the most entropy across positionally indexed sequences from MirGeneDB), yielding 58 unique and most-variant target sequences with conservation levels ranging from 1 to 83 species per sequence (see Appendix C for dataset details). We note that this positional selection assumes curated positional correspondence from the database, which partially reduces the “alignment-free” nature of the approach at the data preparation stage; however, the inference procedure itself operates without computing MSAs. We introduce ConservationWeightedHammingReward (Appendix A.4), which weights target rewards by evolutionary conservation—sequences found in more species receive higher rewards, reflecting purifying selection. To provide informative feedback at each generation step, we also employ ProgressiveHammingReward (Appendix A.5) as the intermediate reward.

For computational efficiency, we restrict the action space to insertions only, reducing the branching factor from *O*(5*n*) to *O*(4*n*) per state while preserving complete sequence generation capability. We intentionally set a training cutoff of 500 iterations (512,000 episodes) instead of training until all target sequences are saturated, simulating an evolutionary search process that focuses on early and mid-stage discovery dynamics rather than merely maximizing target coverage. Under this cutoff, FL-DB achieves coverage of **43/58 (74.1%)** unique let-7 sequences (see Figure 8 for training dynamics). The model preferentially samples conserved sequences, and there is a significant positive correlation between sampling frequency and species count (Spearman *ρ* = 0.509, *p <* 0.001; see Appendix C and Figure 9).

#### Phylogenetic Structure Comparison

To evaluate whether GFlowNet captures evolutionary relationships, we compare traditional phylogenetic reconstruction with our flow-based approach. Figure 3 shows a UPGMA tree constructed from the 43 generated sequences using Hamming distance—a standard approach in molecular phylogenetics. The clustering groups sequences by similarity, with closely related variants sharing evolutionary branches.

**Figure 3:**
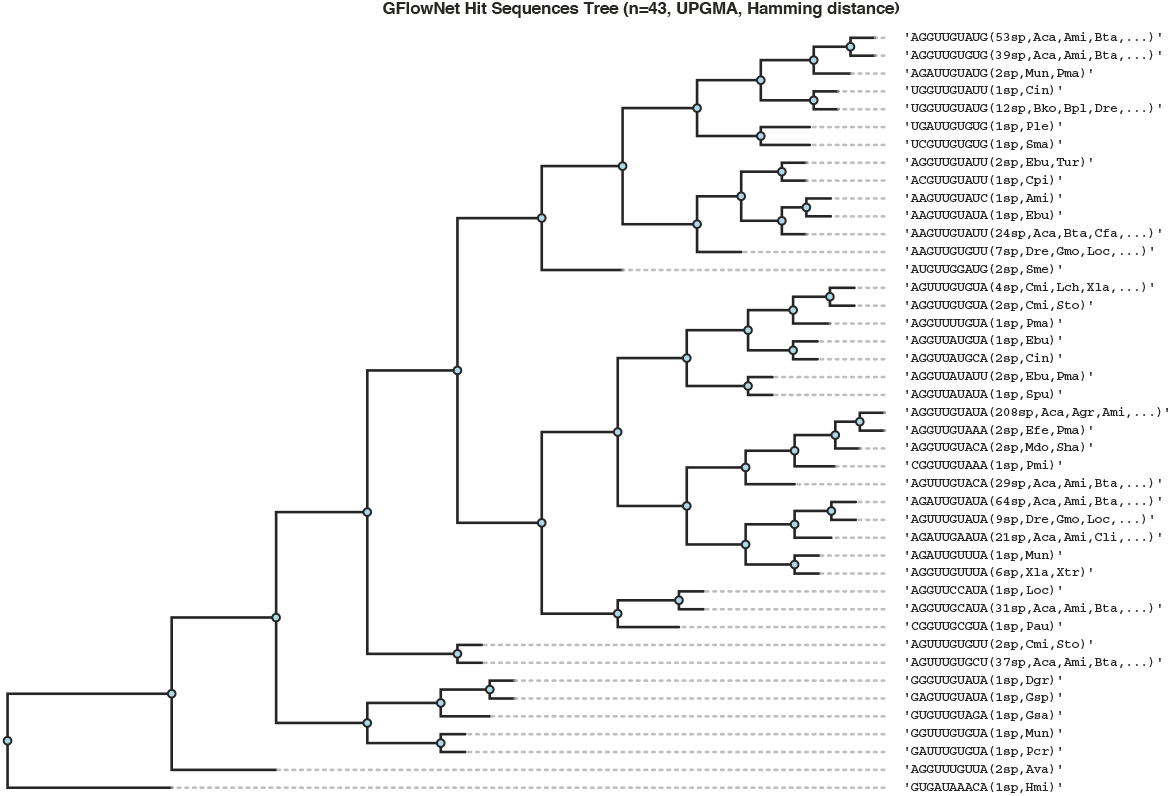
Traditional phylogenetic reconstruction: UPGMA tree of 43 generated let-7 sequences, clustered by Hamming distance. Labels show the sequence and representative species abbreviation. While the tree captures terminal sequence relationships, it does not reveal the generative process or ancestral intermediates.

In contrast, Figure 4 visualizes the state-space DAG constructed from trajectories collected during training, where node size indicates visit frequency and edge width represents flow magnitude. Unlike traditional trees that only capture terminal relationships, the DAG reveals shared intermediate states among related sequences—corresponding to putative common ancestral paths under the generative model. This suggests that even without explicit phylogenetic supervision, the GFlowNet’s learned flow structure qualitatively captures relationships reminiscent of evolutionary branching, providing a complementary view to pairwise distance-based methods. We caution, however, that these inferred “ancestors” reflect the most likely generative prefixes under the learned policy and reward shaping, rather than reconstructions from an explicit evolutionary likelihood model; disentangling reward-induced structure from genuine phylogenetic signal remains an important open question.

**Figure 4:**
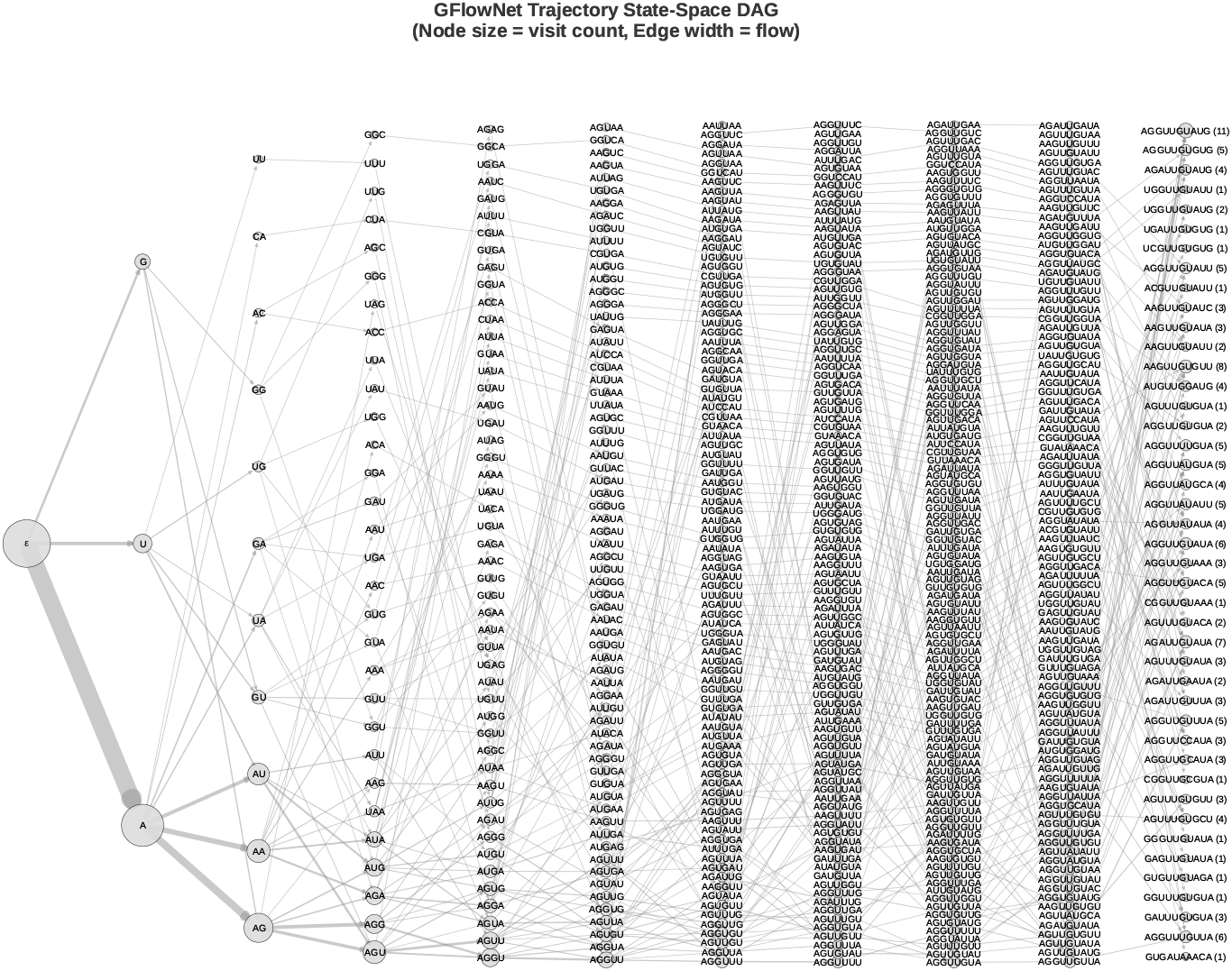
GFlowNet-based evolutionary inference: State-space DAG of generation trajectories for let-7 sequences. Node size indicates visit frequency during sampling; edge width represents flow magnitude. Unlike traditional trees, the DAG reveals shared intermediate states learned by the model, providing a generative view of sequence relationships.

#### Novel Sequence Design

While Figure 4 shows trajectories from training-time stochastic sampling, we can also extract sequences deterministically at inference time. We apply beam search with *k* = 20 to the trained model to find the top-*k* most likely sequences. Figure 5 shows the resulting trajectory DAG, revealing that 5/20 top sequences are known let-7 targets while 15/20 are novel sequences not present in the training set. Notably, these novel sequences cluster near known targets (typically 1–2 Hamming distance), suggesting the model has learned meaningful sequence neighborhoods rather than arbitrary patterns. The shared intermediate states in the beam search DAG provide an alternative view of ancestral relationships: sequences sharing early trajectory prefixes are inferred to have common evolutionary origins. This demonstrates potential for *de novo* sequence design, where trained GFlowNets can propose novel candidates similar to known functional sequences at inference time.

**Figure 5:**
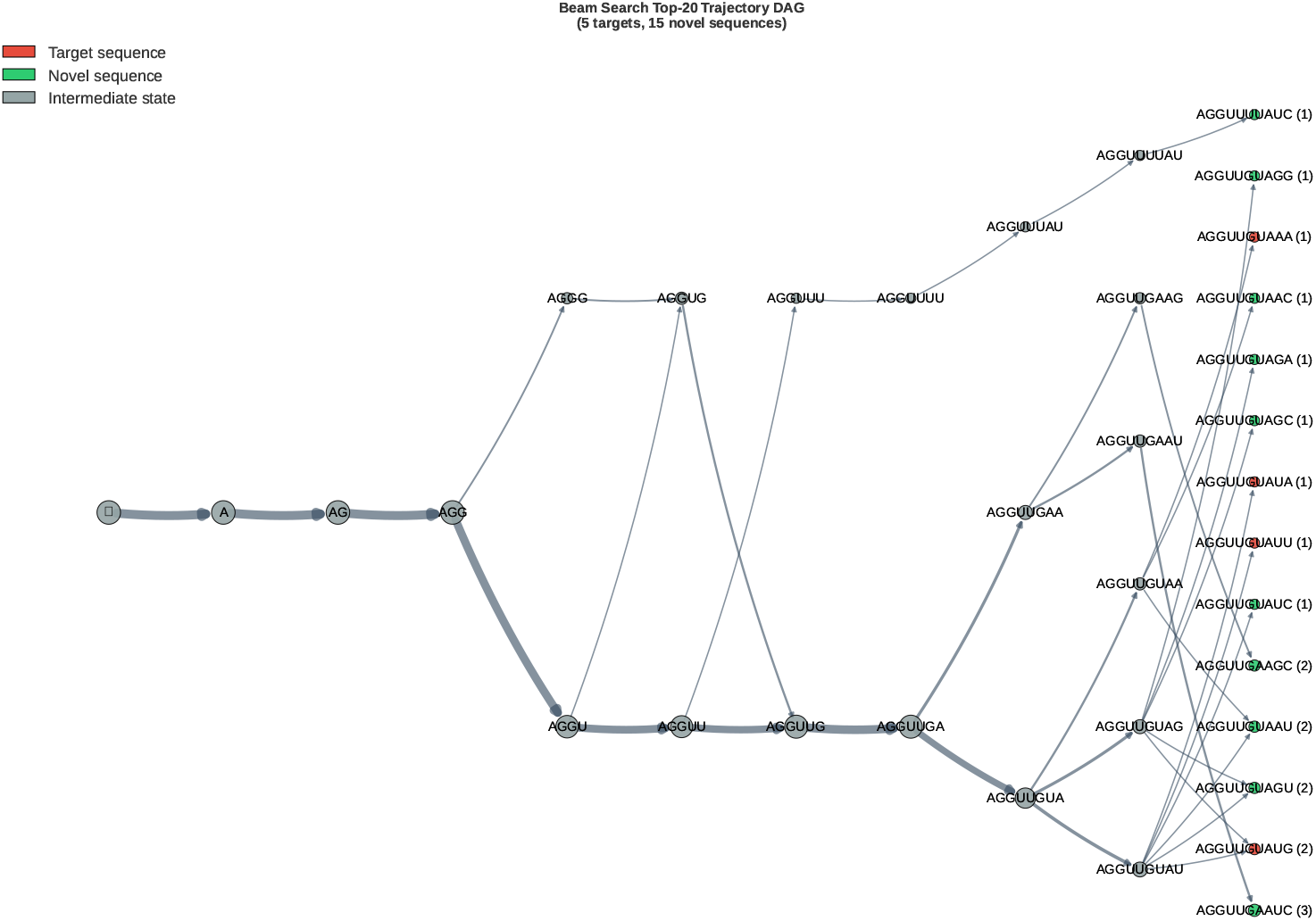
Post-training beam search trajectory DAG for let-7 sequences. Top-20 most likely sequences are extracted via beam search at inference time, with trajectories traced from the root (empty sequence) to terminal states. Red nodes indicate target sequences (5/20), green nodes represent novel sequences (15/20), and gray nodes show shared intermediate states. Novel sequences cluster near known targets (typically 1–2 Hamming distance), suggesting the model learns meaningful sequence neighborhoods.

## 4 Discussion

Our experiments reveal two key insights. First, partial reward signals become increasingly critical as sequence length grows: FL-DB with similarity-based rewards achieves 5× better target coverage than TB on 10bp sequences. This suggests that in sparse reward landscapes, intermediate credit assignment is essential for effective exploration. Second, GFlowNets learn structural relationships among sequences through their flow networks, without explicit phylogenetic supervision. The learned DAG captures branching patterns qualitatively reminiscent of phylogenetic trees, with shared intermediate states suggesting putative common ancestry. Furthermore, beam search at inference time discovers novel sequences near known targets, demonstrating potential for *de novo* sequence design. The ProgressiveHammingReward we introduced addresses weak gradient signals in FL-DB training—a design principle that may generalize to other sequential generative models.

### Limitations

Several limitations warrant discussion. First, our experiments are limited to 10bp sequences; scaling to full-length miRNAs (22bp) or longer sequences remains computationally challenging, and it is unclear whether the observed structural patterns would persist at biologically relevant sequence lengths. Second, our phylogenetic evaluation is entirely qualitative—we lack quantitative comparison with ground-truth phylogenies using standard metrics such as Robinson-Foulds distance or quartet agreement, and have not benchmarked against established tools (e.g., RAxML, MrBayes, IQ-TREE). Third, the inferred “ancestral” states are shaped by the reward function design; the observed branching structure may primarily reflect the geometry induced by the reward rather than evolutionary relationships inherent in the data. Ablations demonstrating stability of the inferred structure across reward variations would strengthen confidence in the phylogenetic interpretation. Fourth, the “alignment-free” framing is partially qualified by the fact that our let-7 data preparation relies on positionally indexed sequences from MirGeneDB, which implicitly assumes some positional correspondence. Finally, evaluation on simulated evolutionary datasets with known true trees and ancestral sequences would provide the most direct validation of whether maximum-flow traceback recovers genuine ancestry rather than reward-induced generative structure.

### Future Directions

Several promising directions emerge from this work. Defining an explicit procedure to extract tree-like objects from the learned trajectory DAG (e.g., a maximum-flow arborescence) would enable quantitative evaluation with standard phylogenetic metrics. Architectural improvements such as hierarchical GFlowNets or attention-based policies may enable scaling to longer sequences. Incorporating phylogenetic likelihood models as reward functions could bridge generative modeling with traditional phylogenetics. Evaluation on simulated datasets with known groundtruth trees, as well as extension to protein sequences and diverse evolutionary datasets, would establish broader applicability.

This work establishes an initial proof-of-concept for alignment-free phylogenetic exploration using generative models, opening new avenues for investigating evolutionary relationships through the lens of learned flow networks.

## Reproducibility Statement

Code for AncestorGFN is available at https://github.com/qhuang20/gflownet-seq-gen. All the data from the experiments is provided, ensuring reproducibility. The miRNA LET-7 family sequences used in this study were obtained from MirGeneDB (https://mirgenedb.org/browse/ALL?family=LET-7&seed=).

## Acknowledgments

We thank Prof. Mathieu Blanchette for advising this project and providing valuable guidance throughout the course of this work. We are grateful to Emily Martin, Dinghuai Zhang, and Emmanuel Bengio for helpful discussions. We also thank the anonymous reviewers for their constructive feedback, which helped improve the clarity and scope of this paper. This work was supported by a research stipend from Prof. Paul François and the Faculty of Medicine ESP-Bioinformatics Scholarship.

## A Reward Function Designs

We implement several reward functions for sequence generation, each addressing different challenges in training GFlowNets.

### A.1 Alignment Reward

The **AlignmentReward** uses Needleman-Wunsch global alignment to provide partial credit:

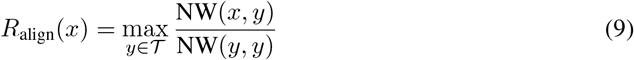

where NW(·,·) is the alignment score (match = +1, mismatch = −1, gap = −1). This is useful when the action space includes insertions and deletions.

### A.3 Entropy-Weighted Hamming Reward

The **EntropyWeightedHammingReward** addresses mode collapse on repetitive sequences by boosting complex (high-entropy) targets:

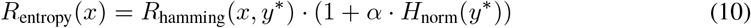

where *H*_norm_(*y*) = − ∑_*c*_ *p*_*c*_ log *p*_*c*_*/* log |A| is the normalized Shannon entropy.

### A.3 Adaptive Hamming Reward

The **AdaptiveHammingReward** dynamically decays rewards for frequently hit targets:

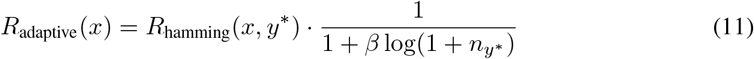

where *n*_*y*_∗ is the cumulative hit count for target *y*^∗^, encouraging exploration of under-sampled targets.

### A.4 Conservation-Weighted Hamming Reward

The **ConservationWeightedHammingReward** weights sequences by evolutionary conservation:

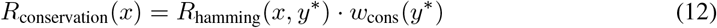

where the conservation weight uses log-scaling:

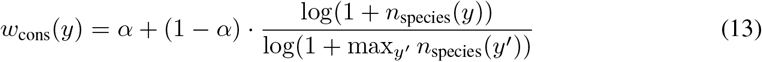

Here *n*_species_(*y*) is the number of species containing sequence *y*, and *α* ensures rare sequences still receive some reward. We use *α* = 0.1, giving conserved sequences (found in ∼80 species) higher reward than singletons.

### A.5 Progressive Hamming Reward

The **ProgressiveHammingReward** improves intermediate reward signals for FL-DB training. Standard Hamming normalization by target length gives weak gradients for partial sequences:

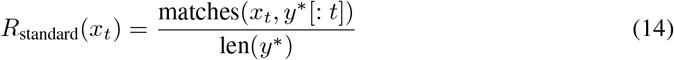

Instead, we normalize by current sequence length for intermediate rewards:

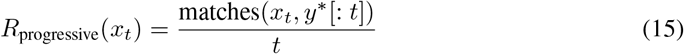

For example, generating towards a 10bp target: at *t* = 1, standard reward gives 1*/*10 = 0.1 (weak), while progressive reward gives 1*/*1 = 1.0 (strong). Terminal rewards still use target length normalization. Figure 6 illustrates this difference.

**Figure 6:**
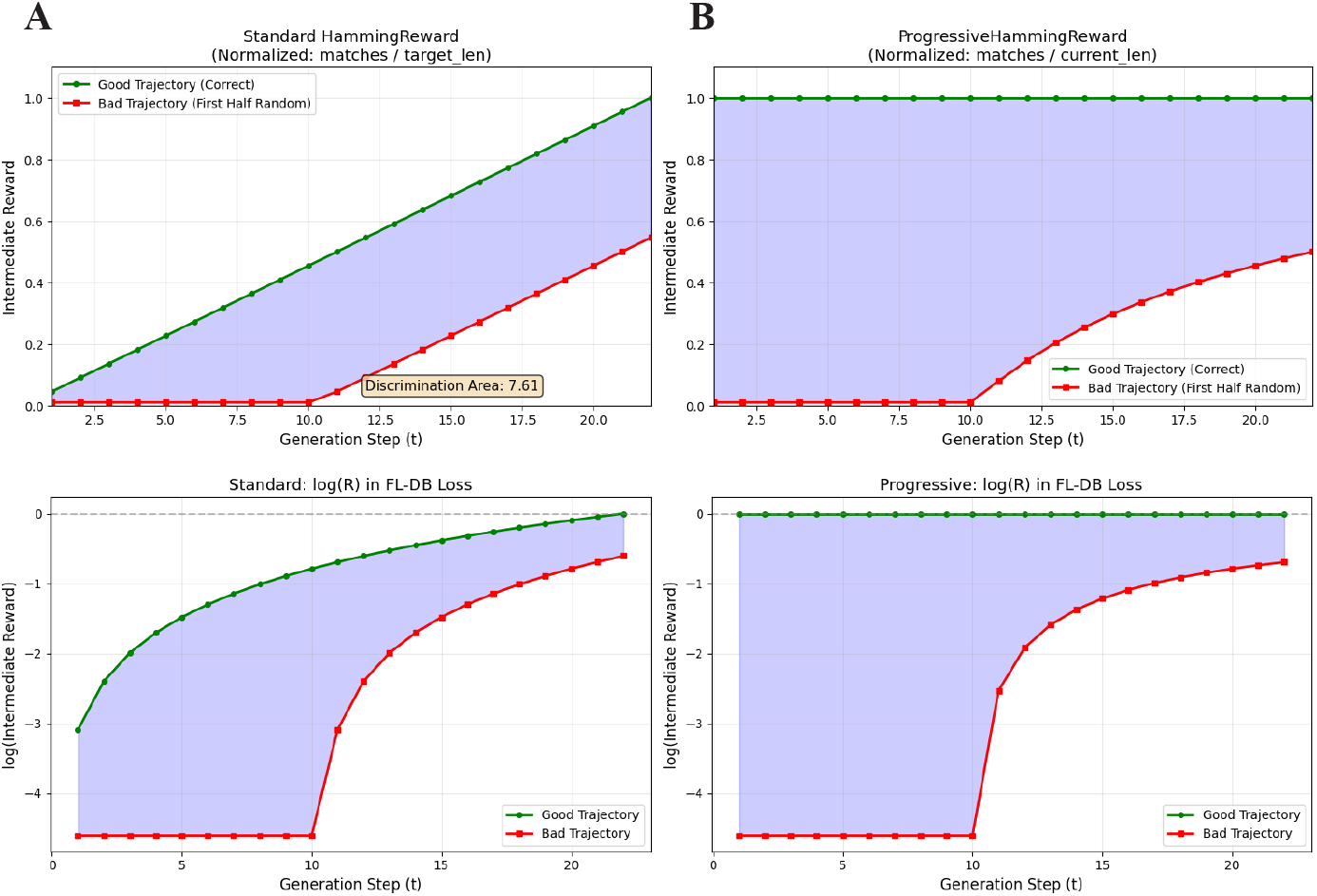
Standard vs. progressive Hamming reward. Progressive normalization by current length provides stronger gradient signals for partial sequences.

## B Training Objective Comparison

Table 1 compares the three GFlowNet training objectives on the 4bp RNA sequence generation task (Case Study 1).

**Table 1:**
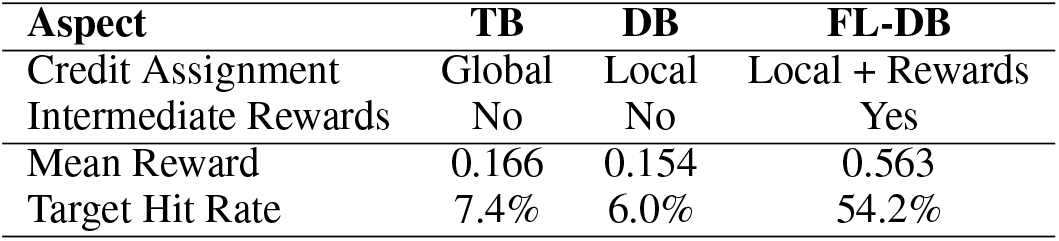
Comparison of GFlowNet training objectives on 4bp sequences. TB/DB use Target-MatchReward (exact match), FL-DB uses AlignmentReward (partial credit).

| Aspect | TB | DB | FL-DB |
| --- | --- | --- | --- |
| Credit Assignment | Global | Local | Local + Rewards |
| Intermediate Rewards | No | No | Yes |
| Mean Reward | 0.166 | 0.154 | 0.563 |
| Target Hit Rate | 7.4% | 6.0% | 54.2% |

**DB** converges faster than TB for simple problems through local flow constraint. **FL-DB** is recommended when meaningful intermediate rewards are available (e.g., partial sequence similarity), achieving significantly higher target hit rates.

## C Let-7 MicroRNA Results

### C.1 Dataset

The let-7 dataset comprises 612 miRNA sequences from 107 species. After extracting positions 10–19 (a 10bp variable region with highest entropy), we obtain 58 unique target sequences with conservation ranging from 1 to 83 species per sequence. Figure 7 shows the distribution of miRNA counts across species (left) and the top conserved sequences ranked by species count (right).

**Figure 7:**
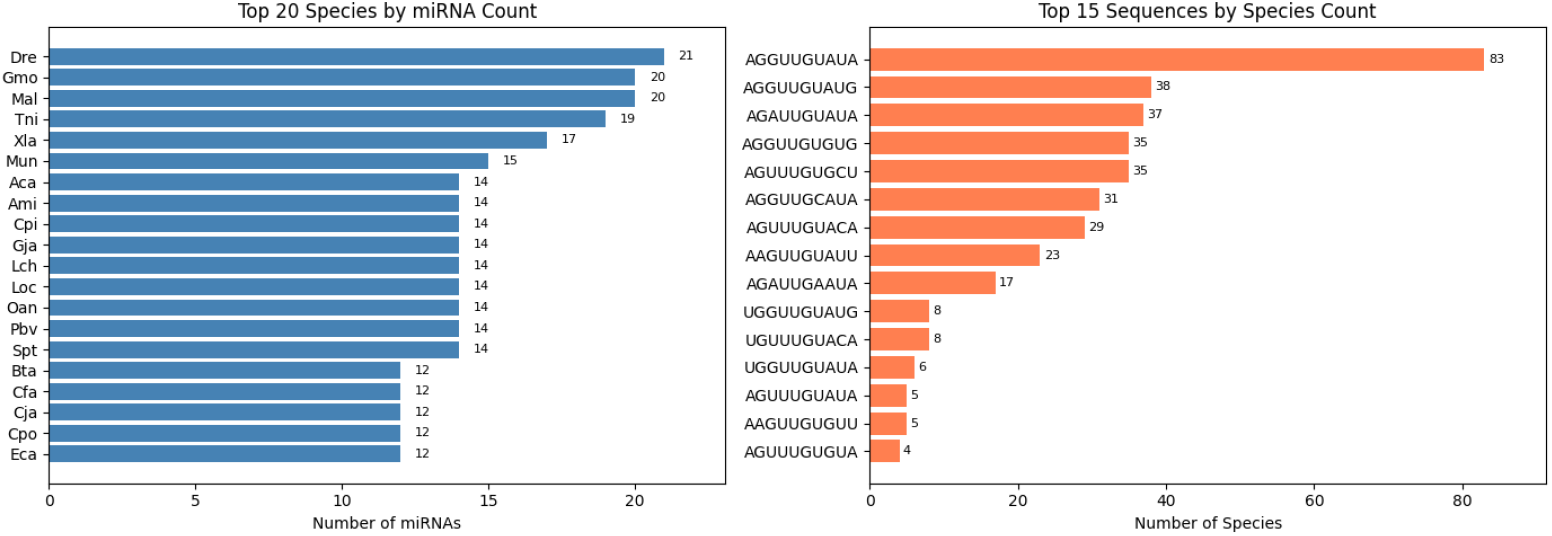
Let-7 dataset statistics. **Left:** Top 20 species by miRNA count. **Right:** Top 15 sequences ranked by conservation (species count).

### C.2 Training Configuration

- **Objective:** FL-DB with ConservationWeightedHammingReward + ProgressiveHammin-gReward
- **Action space:** Insertion-only for efficiency
- **Training:** 500 iterations, batch size 1024, learning rate 3 × 10^−3^
- **Network:** MLP with hidden layers [32, 16, 8]

Figure 8 shows the training dynamics, including loss convergence, log *Z* estimation, target hit rate progression, and cumulative coverage reaching 43/58 (74.1%).

**Figure 8:**
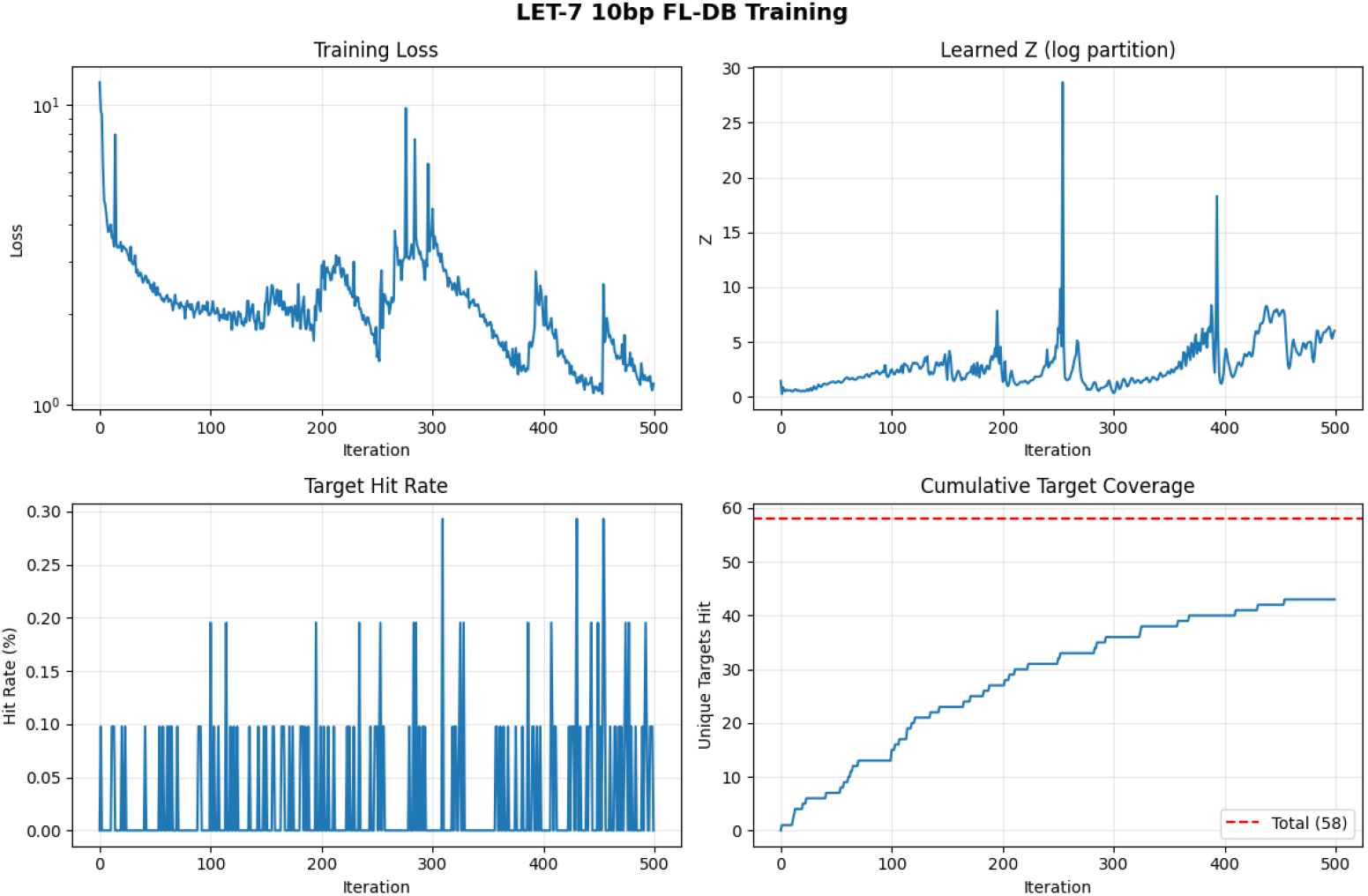
FL-DB training dynamics on let-7 sequences. **Top:** Loss and log *Z* convergence. **Bottom:** Target hit rate and cumulative coverage reaching 43/58 (74.1%).

### C.3 Conservation Correlation

The model preferentially samples conserved sequences (Spearman *ρ* = 0.509, *p <* 0.001), demonstrating that conservation-weighted rewards successfully guide sampling toward evolutionarily important sequences. Figure 9 visualizes this positive correlation between species count and GFlowNet sampling frequency.

**Figure 9:**
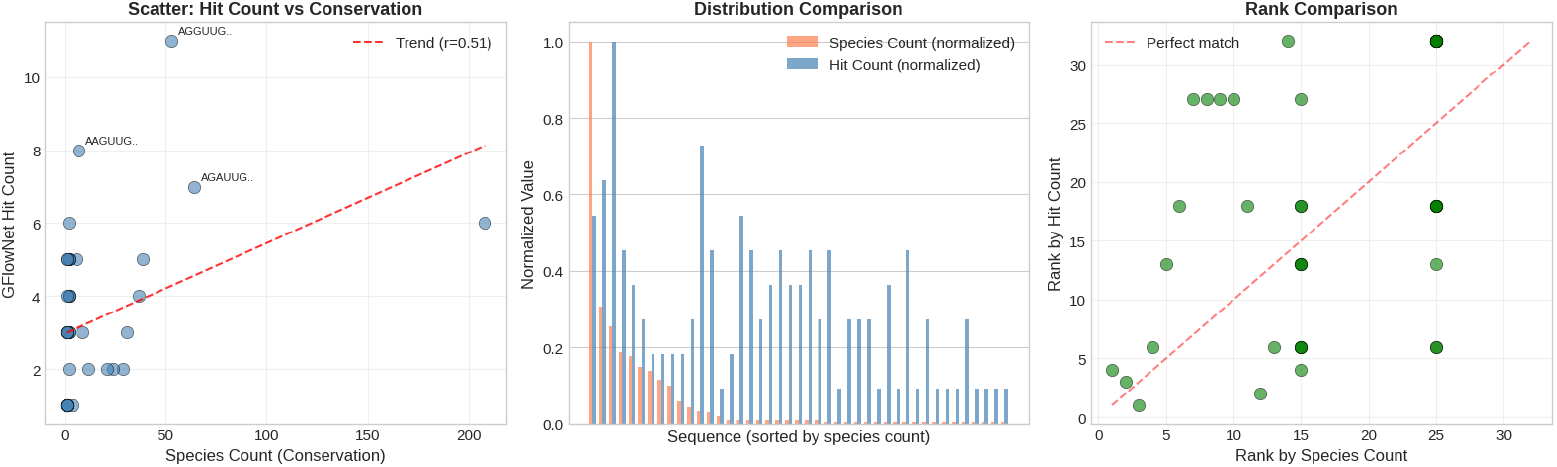
Correlation between species count and GFlowNet sampling frequency.

